# Marker gene fishing for single-cell data with complex heterogeneity

**DOI:** 10.1101/2024.11.03.621735

**Authors:** Yiren Shao, Qi Gao, Liuyang Wang, Dongmei Li, Andrew B. Nixon, Cliburn Chan, Qi-Jing Li, Jichun Xie

**Affiliations:** Department of Data Science, Dana-Farber Cancer Institute, USA; Department of Computational Medicine and Bioinformatics, University of Michigan, USA; Department of Molecular Genetics and Microbiology, Duke University, USA; Department of Clinical and Translational Research, Unversity of Rochester Medical Center, USA; Department of Medicine, Duke University, USA; Department of Biostatistics and Bioinformatics, Duke University, USA; Center for Human Systems Immunology, Duke University, USA; Institute of Molecular and Cell Biology, Agency for Science, Technology and Research, Singapore; Singapore Immunology Network, Agency for Science, Technology and Research, Singapore; Department of Mathematics, Duke University, USA

**Keywords:** Single-cell RNA sequencing, Multi-source heterogeneity, Marker gene identification

## Abstract

In single-cell studies, cells can be characterized with multiple sources of heterogeneity such as cell type, developmental stage, cell cycle phase, activation state, and so on. In some studies, many nuisance sources of heterogeneity (SOH) are of no interest, but may confound the identification of the SOH of interest, and thus affect the accurate annotate the corresponding cell subpopulations. In this paper, we develop B-Lightning, a novel and robust method designed to identify marker genes and cell subpopulations correponding to a SOH (e.g., cell activation status), isolating it from other sources of heterogeneity (e.g., cell type, cell cycle phase). B-Lightning uses an iterative approach to enrich a small set of trustworthy marker genes to more reliable marker genes and boost the signals of the SOH of interest. Multiple numerical and experimental studies showed that B-Lightning outperforms existing methods in terms of sensitivity and robustness in identifying marker genes. Moreover, it increases the power to differentiate cell subpopulations of interest from other heterogeneous cohorts. B-Lightning successfully identified new senescence markers in ciliated cells from human idiopathic pulmonary fibrosis (IPF) lung tissues, new T cell memory and effector markers in the context of SARS-COV-2 infections, and their synchronized patterns which were previously neglected. This paper highlights B-Lightning’s potential as a powerful tool for single-cell data analysis, particularly in complex data sets where sources of heterogeneity of interest are entangled with numerous nuisance factors.

## 1 Introduction

Single-cell sequencing technologies together with computational methods have revolutionized the way we study cellular heterogeneity and gene expression regulation. In many studies, researchers would like to annotate cells and study their multi-aspects of heterogeneity, such as cell types, cell states, and cell-cell interactions.

Two strategies are commonly used to understand a specific source of heterogeneity (SOH). The first strategy is to directly cluster the cells and then characterize the cell clusters^[1,2]^. This strategy highly depends on the clustering quality – inaccurate clustering often leads to misinterpretation of the cell subpopulations. Specifically, the cell subpopulations may just capture the leading and major heterogeneity source, such as cell types. If one would like to focus on a minor SOH with weak signal-to-noise ratio such as senescence, this strategy often fails to separate the senescent cells. The second strategy is to identify marker genes for the SOH of interest and then use these marker genes to annotate the corresponding cell subpopulations. This strategy is more direct and interpretable, but it requires many trustworthy marker genes. Often, if the number of trustworthy marker genes is limited, they may not be sufficient to differentiate the cell subpopulations of interest.

One way to address this issue is to start with a few trustworthy marker genes for the SOH of interest, and then use them as “bait” to fish out more marker genes for the target SOH. Increasing the number of marker genes will help boost the signal-to-noise ratio and better differentiate the corresponding cell subpopulations of interest.

To the best of our knowledge, two approaches are available for fishing out new genes. An ad-hoc approach is to derive cellular gene set enrichment scores based on the known marker genes, use these scores to group cells into marker-enriched and non-enriched subpopulations and finally perform differential gene expression (DGE) analysis on these two subpopulations to find new marker genes. This approach uses the same dataset in clustering and DGE analysis, and thus inflates type I error due to “double-dipping”^[3,4]^. The other approach, GeneFishing^[5]^, partitions the genes many times and checks for a consistency trend of gene clusters to identify new marker genes. However, the gene clustering is based on dimension reduction on the gene expression matrix. When the UMI counts of scRNA-seq are sparse, informative dimension reduction on its co-expression matrix is difficult to achieve, which can affect GeneFishing’s downstream performance.

In this paper, we introduce a new method, B-Lightning, that iterates between identifying new marker genes and extracting the corresponding cell subpopulations. The key idea is to use an iterative procedure to gradually expand the feature gene set: at each step we find the candidates using DGE analysis and then use the connectivity of the gene co-expression network to filter out the false discoveries. This idea is based on our previous work showing that marker genes from the same SOH tend to exhibit high connectivity in the gene co-expression network^[6]^. After using the connectivity threshold to filter candidate marker genes, we update the cellular feature score (CFS) and the process is repeated until convergence. The workflow of B-Lightning is shown in Figure 1.

**Figure 1:**
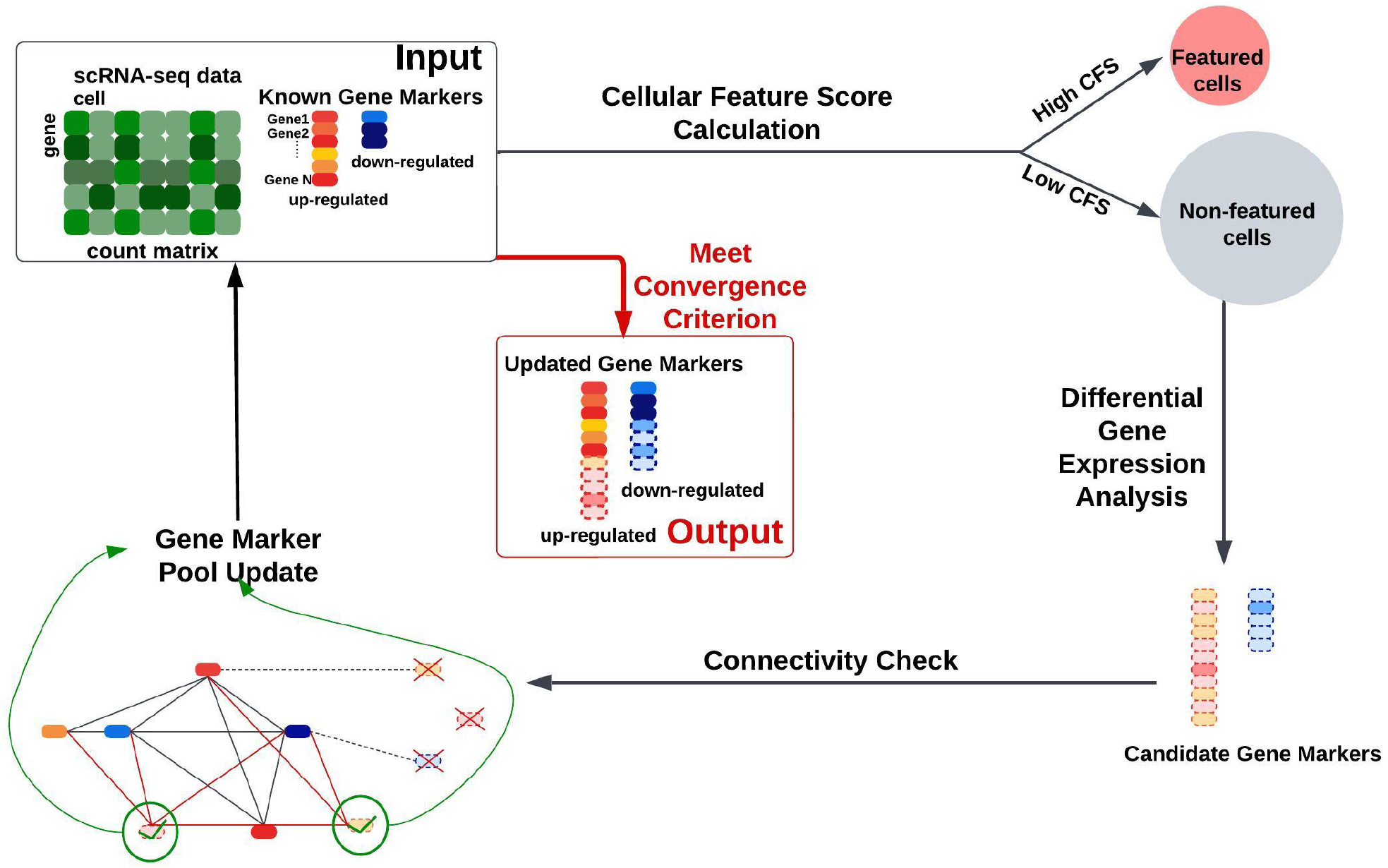
Workflow of B-Lightning. Known context-specific markers are used as input markers to define a preliminary cellular feature score (CFS) for each cell. Cells with high CFS will be tagged as featured cells. Then, DGE analysis is performed on CFS between feature and non-feature cells to select new candidate markers. Those candidate markers showing high connectivity with the initial input markers are identified as the new markers; they will be used as the input of the next iteration to re-calculate CFS and update the candidate markers. The algorithm is considered to have converged when there is no update in the two previous iterations.

**Figure 2:**
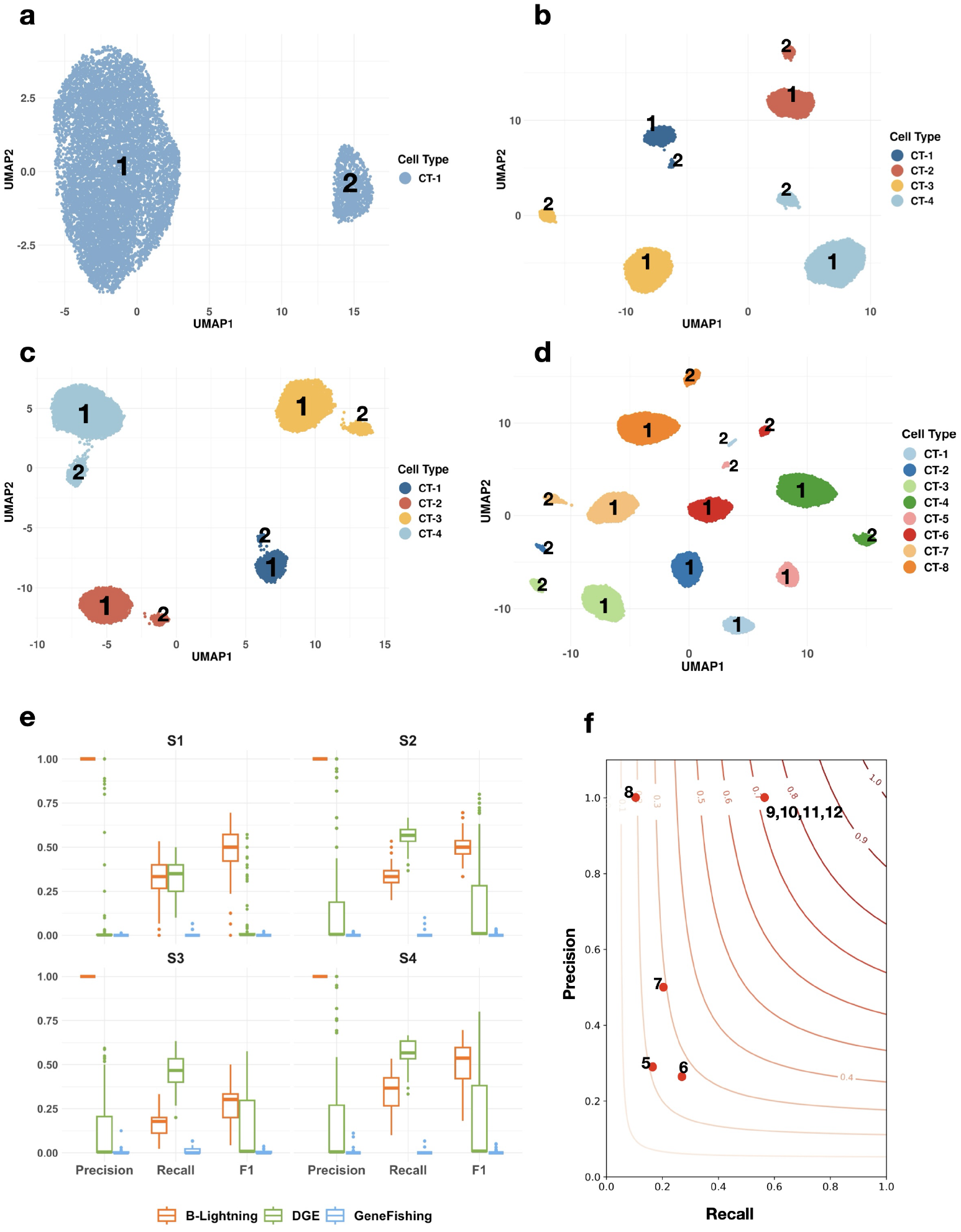
Comparison of methods on simulated data. **(a-d) UMAPs of simulated datasets under settings S1-S4**. We randomly selected a simulated dataset from each setting and show their gene expression UMAPs. Each cell was colored by its cell type and labeled by its cell state (1 for State 1, and 2 for State 2). **(e) Performance comparison between B-Lightning, DGE analysis, and GeneFishing**. Boxplots of precision, recall and F1 score on identification of cell state feature genes of 3 methods: B-Lightning(orange), DEG analysis(green), GeneFishing(blue). Performance was evaluated based on 100 replications of each simulated dataset. **(f) Performance of B-Lightning with different number of inputs**. Contour plot with precision (y-axis), recall (x-axis) and F1 score (contour line) on identification of cell state feature genes via B-Lightning in S1. The number by each dot refers to the quantity of input genes.

We have compared the performance of B-Lightning and other methods via simulated and experimental data analysis. The results show that B-Lightning is more sensitive and robust than competing methods. It successfully identified new senescence markers for ciliated cells in human Idiopathic Pulmonary Fibrosis (IPF) lung data samples. Furthermore, it identified new T cell memory and effector makers, and revealed a previously neglected synchronized pattern in the transformation of effective and memory T cells during SARS-COV-2 infections.

## Methods

### B-Lightning algorithm

B-Lightning is an iterative algorithm alternating between defining cellular feature scores (CFS) and identifying new feature genes. It starts from a set of known feature genes, and aims to iteratively expand the set of feature genes and to identify the cell subpopulation corresponding to these feature genes. Algorithm 1 lists its main steps, and the details of each step is described below.

#### Algorithm 1

B-Lightning algorithm.

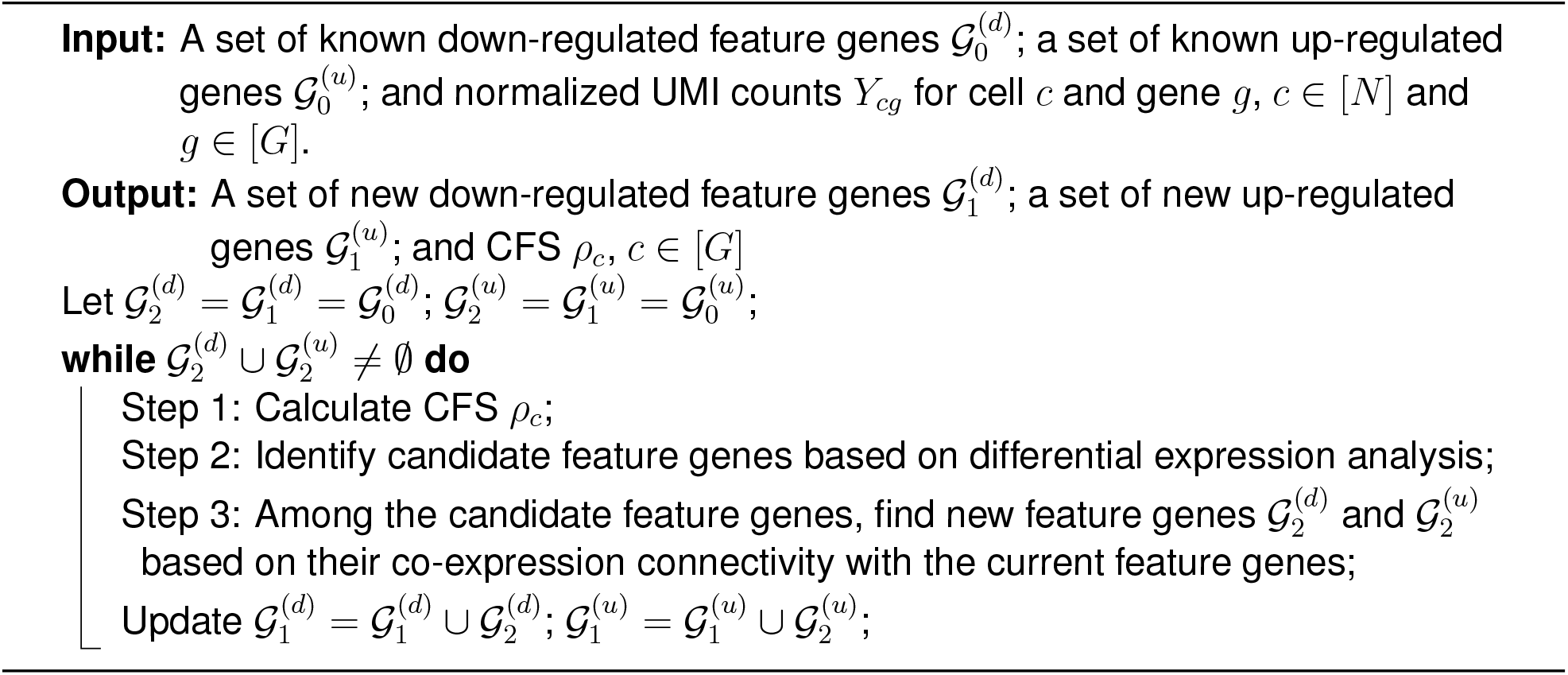

#### Step 1: Calculate CFS

The CFS is calculated based on the current up-regulated feature gene set 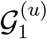 and the down-regulated feature gene set 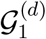. Let *Y*_*cg*_ be the normalized UMI for cell *c* and gene *g*. For any 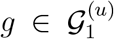, the up-regulated gene-specific CFS is 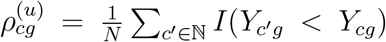; for any 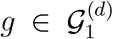, the down-regulated gene-specific CFS is 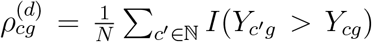. These are the sample tail probabilities of gene *g* being over-expressed or under-expressed in cell *c*. Then, the CFS for cell *c* is defined as

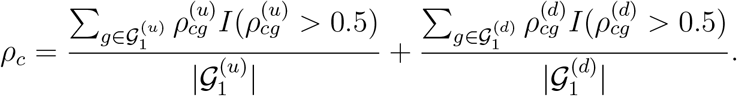

#### Step 2: Define candidate feature genes

Based on CFS, we divide all *N* cells into two groups, featured cells and non-featured cells. The default cut-off for cell grouping is 90% quantile of CFS. Cells with higher CFS are identified as featured cells. Then, we identify new gene markers by performing differential gene expression (DGE) analysis for the two groups. By default, we used MAST^[7]^, but in principle other DGE method should also work. False discovery rate (FDR) is controlled by the Benjamini and Hochberg (BH)^[8]^ procedure at 5%.

#### Step 3: Find new feature genes

For any initial input gene *i* and candidate gene *j*, we used quantile associations *S*_*ij*_ to measure their co-expression. Then we draw highly variable genes, and compute their quantile associations *S*_*ik*_ with the initial input gene *i* too. These *S*_*ik*_ form a benchmark null distribution for *S*_*ij*_. Gene *i* and gene *j* are connected if *S*_*ij*_ is above the 95% quantiles of all the *S*_*ik*_ (Supplementary Note 1). Previous work^[6]^ has shown that genes within the same gene set tend to co-express. Therefore, we use this property to filter out feature genes. Only new gene markers connected with most of the initial input genes will be added to the feature gene sets 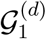 and 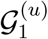.

## Competing Methods

### DGE analysis

We used the MAST^[7]^ procedure, a hurdle model-based DGE analysis method that models the modified gene set variation analysis (GSVA) score^[9]^. The modified GSVA score was calculated by deducting the GSVA scores of the down-regulated gene marker set from the GSVA score of the up-regulated gene marker set. Genes were identified as markers if they were significant at a FDR less than 5% and with a log2 fold change greater than 0.025.

### GeneFishing

GeneFishing^[5]^ is a computational method designed to use “bait” genes for “fishing out” new genes based on bulk RNA-seq data. We used the same bait genes as B-Lightning and repeated 100 rounds of spectral clustering to compute the capture frequency rates (CFRs) for each candidate gene. We identified genes with CFR > 0.9 as gene markers, as recommended in the GeneFishing paper.

### Simulated data generation and analysis

We simulated four scenarios (dataset S1-S4) using the R package GeneScape^[10]^. GeneScape allows users to simulate scRNA-seq data with complex heterogeneity structures. Here, we consider the cases when cells have both cell type and cell state heterogeneities. S1 contains 10000 cells of the same cell type and 5000 genes. Among all the cells, 10% cells have a differential cell state with 30 genes differentially expressed. We call these cells state feature cells, and these genes state feature genes. S2 and S3 contain 10000 cells of 4 cell types, with 1000, 2000, 3000, and 4000 cells in each cell type, respectively. Cell types are defined by 360 cell type feature genes. Within each cell type, 10% are state feature cells, marked by 30 state feature genes. Specifically, in S2, feature cells of different cell types share the same state feature genes; while in S3, featured cells of different cell types shared only 15 feature genes only. S4 contains 5000 genes and 20000 cells of 8 cell types, two have 1000, 2000, 3000, and 4000 cells, respectively. For each cell type, 10% cells were featured cells with 30 shared feature genes across all cell types.

In each scenario, we used 100 different seeds to generate 100 datasets. Then we pre-processed all the datasets by removing cells with less than 5% of the genes expressed and genes with expression in less than 5% cells. After using NormalizeData function in Seurat to normalize the data follwed by log2 transformation, we applied B-Lightning, DGE analysis, and GeneFishing to each dataset. For B-Lighting and GeneFishing, we input 10 state feature genes as “bait” genes. We used 90% quantile of cellular feature scores to group cells.

## Experimental data analysis

### IPF2 and IPF1

These two datasets were collected from the existing study on idiopathic pulmonary fibrosis (IPF)^[11]^.

IPF1 contains the pre-annotated “KRT5-/KRT17+ cells” and “proliferating epithelial cells”. “KRT5-/KRT17+ cells” were treated as senescent cells as in the original paper of the study while “proliferating epithelial cells” were treated as non-senescent cells.

IPF2 contains the pre-annotated “ciliated cells” from IPF patients. We preprocessed IPF2 to filter out cells with less than 5% of the genes expressed and genes expressed in less than 5% of the cells. Then we used NormalizeData function in Seurat to normalize the data before the log2 transformation.

To obtain the bait genes, we initiated from the SenMayo^[12]^, a recently published senescence marker set. Previous studies have shown that senescence marker exhibit tissue-specific and cell-type-specific profiles^[13,14]^. To make sure the identified senescence markers are specific to ciliated cells, we performed DGE analysis on these genes on the “KRT5-/KRT17+ cells” and “proliferating epithelial cells” in IPF1. Those genes with FDR < 0.05 and absolute value of log fold change > 0.25 were selected as bait genes. Among all bait genes, 20 are up-regulated, and 2 are down-regulated (Supplementary Table 1).

When applied B-Lightning on IPF2, we used the 90% quantile of CFS as cutoff to label senescent cells. At each iteration, we requested the newly identified senescence markers to connect with more than 20% of the known gene markers. In total, we found 54 new senescence markers. We then checked the log fold changes and P values of these markers between the KRT5-/KRT17+ cells (senescent cells) and proliferating epithelial cells (non-senescent cells) in IPF1. Among all the new senescence markers, 11 markers have significant P values and log fold change at the same direction: 7 are up-regulated, and 4 are down-regulated. (Figure 3b).

**Figure 3:**
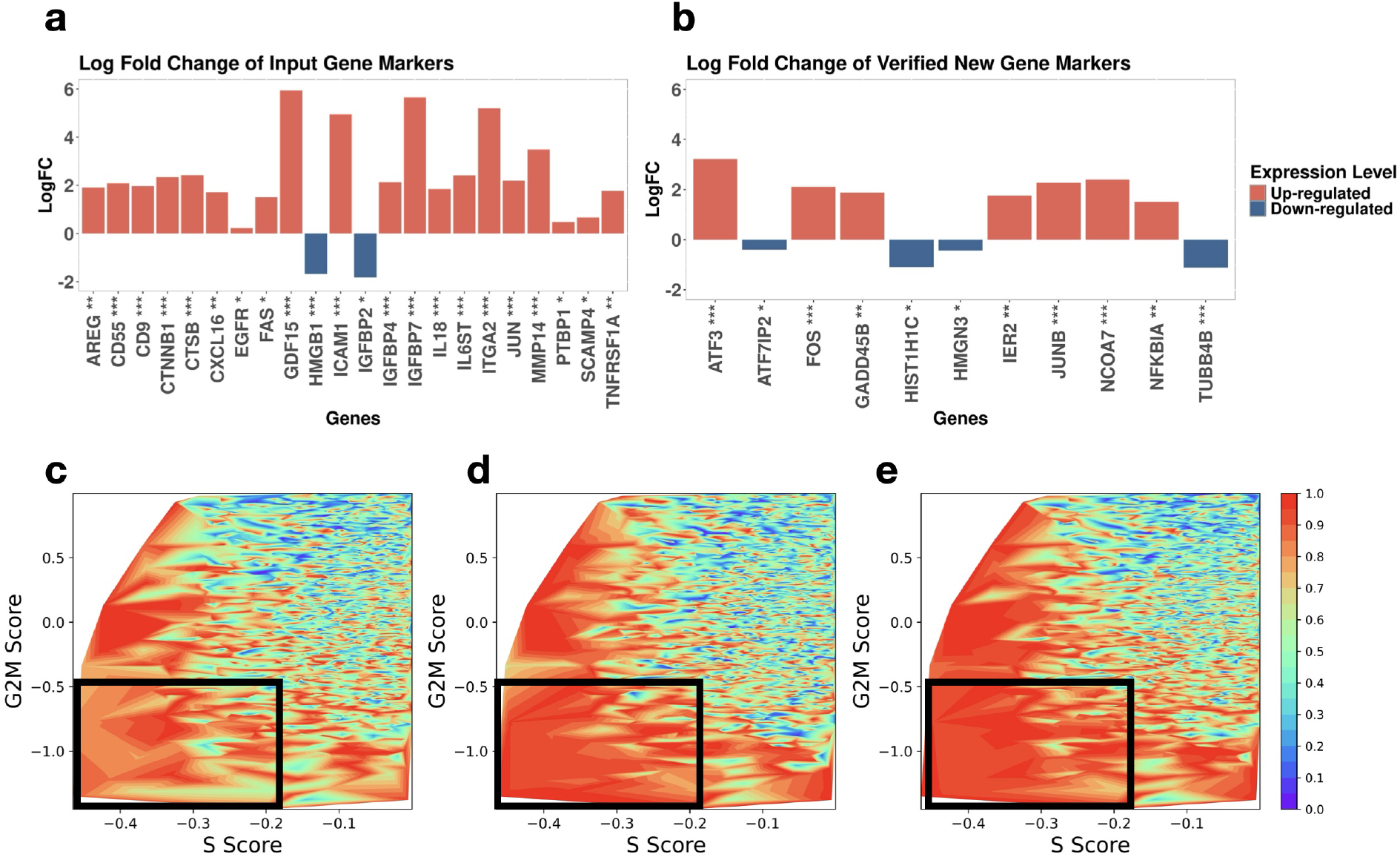
B-Lightning identified new senescence markers for lung ciliated cells. (a-b) Log fold change of genes. (a) shows the log fold change of input genes and their p-values between the KRT5-/KRT17+ cells and proliferating epithelial cells in IPF1. Then we applied B-Lightning to IPF2 to identify new senescence markers. Next, we checked the log fold change of newly identified markers and their p-values between the KRT5-/KRT17+ cells and proliferating epithelial cells in IPF1. (b) shows the 11 genes having significant p-values. Asterisks represent the significance level of the difference between senescent and non-senescent cells, ∗: p-value ≤ 10^−2^, ∗∗: p-value ≤ 10^−4^, ∗ ∗ ∗: p-value ≤ 10^− 6^. **(c-d) B-Lightning enriched ciliated senescent cells**. We calculated CFS by (c) the input genes, (d) the significant new genes, and (e) the combination of input genes and significant new genes. We also plotted the cell cycle S and G2M scores for each cell. Contour plots with S score (x-axis), and G2M score (y-axis), colored by each cell’s percentile rank of CFS were plotted. Only those cells with G2M Score < 1 and S Score < 0 were included. After B-Lightning enrichment, the cells in the low S and G2M score region (in the black rectangles) are more highly ranked in senescence CFS scores.

Next, we calculated the senescence CFS based on three gene sets, the bait genes, the 11 new significant senescence markers, and the combined markers, leading to three different sets of CFS. For each set of CFS, we normalized the CFS for each cell based on its percentile among all cells. We also used Seurat V3^[1]^ function CellCycleScoring to calculate cell cycle scores (G2M score and S score). Cells receiving higher positive G2M (or S) scores are more likely to be cells in the G2M (or S) phase. The cellular scores are then compared with the three sets of the CFS to see whether the new senescence markers help to improve the senescent cell calling.

### 74-year-old Female COVID-19 Patient Airway CD4/CD8 (74y-COVID)

This dataset contains longitudinal single-cell RNA-seq data from a 74-year-old female patient with severe COVID-19. CD4 and CD8 T cells were collected from the airway wash 1 day, 3 days, 4 days, 5 days, 6 days, 7 days and 8 days post-intubation^[15]^.

We pre-processed the datasets to screen out cells with higher than 20% of mitochondrial reads, molecular counts more than 30000, cellular UMI counts less than 100 or more than 6000, and genes expressed in less than 1% of the cells. Then we used the NormalizeData function in Seurat V3 to normalize the data before the log2 transformation. We used the 90% quantile of CFS as cutoff to group cells into senescent cells and non-senescent cells. Newly detected gene markers were required to be connected with more than 30% of the input gene markers for CD4 cells and more than 40% of the input gene markers for CD8 cells.

Published lineage markers (Supplementary Table 3) of CD4/CD8 effector/memory cells were used as the input markers to fish out new CD4/CD8 effector/memory markers. After applying B-Lightning, we identified 13 new effector and 46 new memory markers for CD4 cells, and 32 new effector and 32 new memory markers for CD8 cells (Supplementary Table 4). The input markers and the newly identified markers were aggregated to form the combined markers. Four CFS scores were calculated for each cell: the memory CFS and effector CFS based on the input markers and the memory CFS and effector CFS based on the combined markers. Then, we plotted the density distributions of the memory CFS (Figure 4a, Supplementary Figure 3a, Supplementary Figure 5a, c) and effector CFS for the two sets of markers (Figure 4b, Supplementary Figure 3b, Supplementary Figure 5b, d).

**Figure 4:**
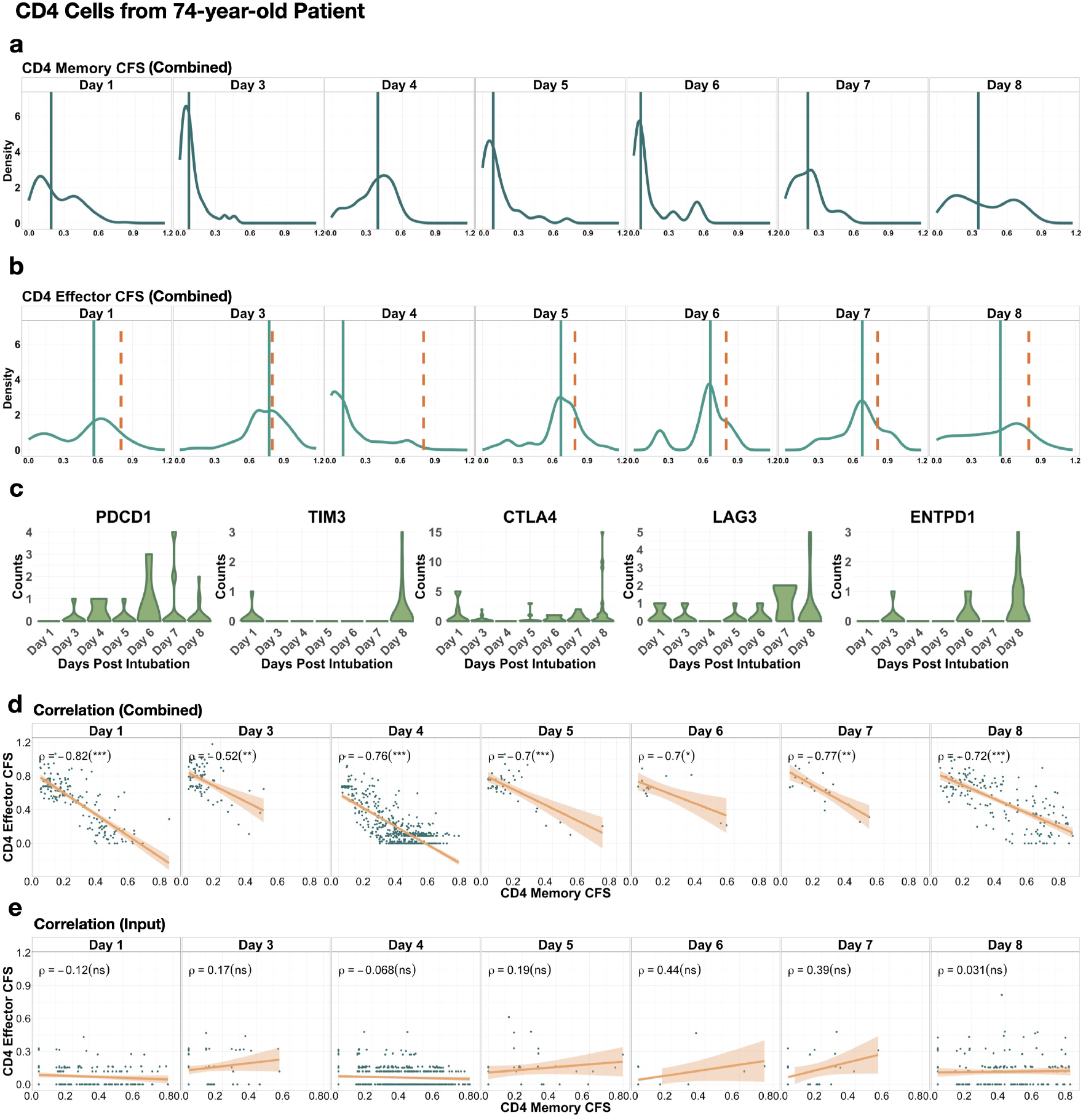
B-Lightning identified new T effector and memory markers in COVID-19 patients and revealed synchronized dynamics between effector and memory T cells. (a-b) Density plots of CD4 CFS. (a) The CD4 memory CFS. (b) The CD4 effector CFS. CFS was calculated by the combined markers. The vertical solid line refers to the median of the combined memory marker CFS. The orange dashed line refers to the combined effector marker CFS threshold (0.8) used to select cells with high effector CFS. **(c) Expression counts of CD4 cell exhaustion markers**. Violin plots of T cell exhaustion markers PDCD1, TIM3, CTLA4, LAG3 and ENTP1 (CD39) of effector CD4 T cells (with combined marker effector CFS > 0.8) on days post intubation. **(d-e) Scatter Plots of CD4 memory CFS vs. effector CFS**. (d) The combined marker memory CFS is the x-axis, and the combined marker effector CFS is the y-axis. (e) The input marker memory CFS is the x-axis, and the input marker effector CFS is the y-axis. *ρ* stands for the Pearson correlation between the memory and effector markers. Asterisks/ns in the parentheses represent the significance level of the correlation between senescent and non-senescent cells, ∗: p-value ≤ 10^−2^, ∗∗: p-value ≤ 10^−4^, ∗ ∗ ∗: p-value ≤ 10^−6^, ns: non-significant. Each subplot extracted the cells collected on a specific day after intubation.

### 49-year-old Male and 66-year-old Female COVID-19 Patients Airway CD4/CD8 (49y-COVID, 66y-COVID)

This dataset contains longitudinal single-cell RNA-seq data from a 49-year-old male patient and a 66-year-old female patient with severe COVID-19. CD4 and CD8 T cells were collected from the airway wash 1 day, 2 days, 3 days post-intubation and 2 days, 3 days, 4 days 7 days post-intubation^[15]^ respectively. We pre-processed the datasets to screen out cells with higher than 20% of mitochondrial reads, molecular counts more than 30000, cellular UMI counts less than 100 or more than 6000, and genes expressed in less than 1% of the cells. Then we used the NormalizeData function in Seurat V3 to normalize the data before the log2 transformation. We included CD4 and CD8 cells based on the cell type annotations in the original paper^[15]^. Four CFS scores were calculated for each cell: the memory CFS and effector CFS based on the input markers and the memory CFS and effector CFS based on the combined markers identified using the 74y-COVID dataset (Figure 5a, b, e, f, Supplementary Figure 4b, f, Supplementary Figure 6).

**Figure 5:**
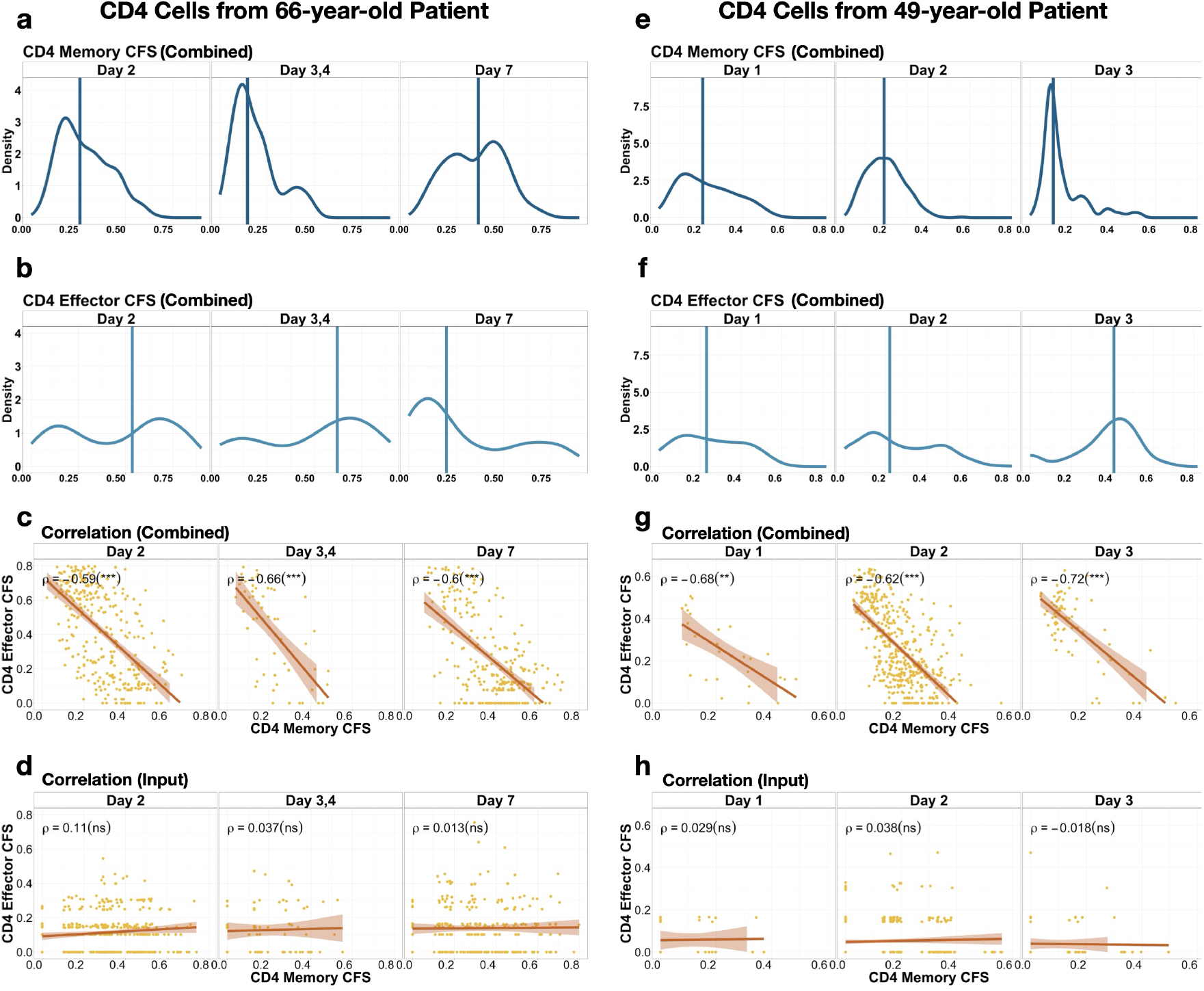
The synchronized dynamics between effector and memory T cells are consistent across different COVID-19 patients. (a-b, e-f) Density plots of combined marker CFS for CD4 cells from the 66-year-old and 49-year-old Patients. (a,e) The combined marker CD4 memory CFS. (b,f) The combined marker CD4 effector CFS. The vertical solid line refers to the median of the combined marker effector CFS. **(c-d, g-h) Scatter Plots of CD4 memory CFS vs. effector CFS**. (c,g) The combined marker memory CFS is the x-axis, and the combined marker effector CFS is the y-axis. (d,h) The input marker memory CFS is the x-axis, and the input marker effector CFS is the y-axis. *ρ* stands for the Pearson correlation between the memory and effector markers. Asterisks/ns in the parentheses represent the significance level of the correlation between senescent and non-senescent cells, ∗: p-value ≤ 10^−2^, ∗∗: p-value ≤ 10^−4^, ∗ ∗ ∗: p-value ≤ 10^−6^, ns: non-significant. Each subplot extracted the cells collected on a specific day after intubation.

## Results

To examine the performance of B-Lightning, we applied it to the simulated datasets (S1-S4) and compare the results with two competing methods, DGE analysis and GeneFishing. We also applied B-Lightning to two experimental studies, a senescence study in IPF ciliated cells and a T cell study in cells from subjects with COVID-19. The competing methods and experimental datasets are described in Methods.

### B-Lightning “fished out” new cell state marker genes in data with complex heterogeneity

We systematically compared the performance of B-Lightning with two competing methods, DEG analysis and GeneFishing, in a numerical study. We simulated scRNA-seq data with SOH from only cell state (S1, Figure 2a), and from cell types and cell states (S2-S4, Figures 2b-d): the latter three datasets contain two orthogonal SOH: cell types and cell states. “Cell state” is the SOH, with lower signal-to-noise ratio compared with “cell type”, considered as the nuisance SOH in this study. S1 consists of 1 cell type, S2 and S3 consist of 4 cell types, and S4 consists of 8 cell types. Each cell types have their corresponding up-regulated and down-regulated feature genes. In terms of cell states, all datasets have 2 cell states. Under S1, S2, and S4, the cell states across different cell types share the same state feature genes. Under S3, the cell state feature genes differ across different cell types; only 50% of state feature genes are shared across different cell types. Thus, S2-S4 are more challenging cases than S1.

Our goal is to use a few established cell state specific “bait genes” to “fish out” new cell state markers. We simulated 100 datasets under each setting (S1-S4) to compare B-Lightning and with the standard DEG analysis and GeneFishing. The performance of each method was evaluated using the precision, recall, and the F1 score of the newly identified state feature genes (Figure 2e). Here, the F1 score is the harmonic mean of precision and recall, representing the balance between the two.

Under all settings, B-Lightning has the high median precision among all methods. In fact, in over 95% of all simulated datasets, B-Lightning has precision equal to 1, indicating that all of its identified cell state markers are true. In contrast, DGE has a large variation in precision, suggesting its lack of robustness. GeneFishing, has consistently low precision, suggesting many of its identified genes are false discoveries. In terms of recall, although DGE has the highest median recall under S2-S4, its precision is highly inflated. GeneFishing has poor recall, probably because it is not designed to fish out feature genes for single-cell data with confounding heterogeneity SOH. Overall, B-Lightning has the best F1 score in identifying new state feature genes: it has the highest median F1 score under all settings, as well as a small variance across 100 repetitions.

The main reason that B-Lightning outperforms DGE and GeneFishing is that B-Lightning uses both the marginal gene expression and connectivity to identify feature genes from the same SOH^[6]^. Considering sparsity and noisiness in the single-cell data, single gene marginal expression or single graph edge connectivity is not sufficient to screen out marker genes. However, checking connectivity over multiple edges can increase the accuracy and robustness of the screening procedure and improve performance.

To further verify the robustness of B-Lightning, we applied it with different number of input markers. For a randomly simulated dataset under S1, as the number of input gene markers increases till 9, B-Lightning’s F1 score increases; after that, adding more input markers does not further improve the performance of B-Lightning (Figure 2f). The precision and recall are the same as we change the number of input markers from 9 to 12. The results demonstrate the robustness of B-Lightning to the input markers.

### B-Lightning identified new potential senescence markers in lung ciliated cells

Cellular senescence and lung fibrosis are strongly associated^[16,17]^; thus, we employed B-Lightning to identify senescence gene markers in cells from lung tissues of subjects with idiopathic pulmonary fibrosis (IPF). An existing dataset, IPF1, contains ciliated cells that have been annotated as KRT5-/KRT17+ cells (considered as a senescence enriched cell population) and proliferating epithelial cells (non-senescent cells). To select some high-confidence senescence input markers, we used the intersection gene set of a recently published senescence gene panel, SenMayo^[12]^, and the DGE genes in the IPF1 dataset (Figure 3a, Methods). This is to ensure the input senescence markers fit the IPF lung ciliated cell context. There are 20 upregulated and 2 down-regulated input gene markers. These genes’ log fold change between the KRT5-/KRT17+ cells and proliferating epithelial cells in IPF1, as well as the corresponding p-values, are shown in Figure 3a.

After we obtained a high-confidence input gene set, we applied B-Lightning to IPF2, another independent lung IPF single-cell dataset, to identify new senescence markers. B-Lightning identified additional 13 up-regulated and 41 down-regulated genes (Supplementary Table 1). Then we went back to IPF1 to check the log-odds ratio and the p-values between the KRT5-/KRT17+ cells (senescent cells) and proliferating epithelial cells (non-senescent cells). We found that 11 newly identified senescence markers have the same direction of log fold change in IPF1 as in IPF2, as well as significant p-values (Figure 3b). Among the 11 newly identified genes, nine of them have direct connections with senescence based on previous studies (Supplementary Table 2). The remaining two markers, *TUBB4* and *HMGN3*, play important roles in cell division and response to cellular stress such as DNA damage; thus may also play a role in senescence (Supplementary Table 2).

We then compared the three sets of cellular feature scores (CFS) calculated based on the input genes, the 11 new significant senescence markers, and the combined markers. A good set of CFS score should indicate cellular senescence so that cells with high CFS are likely to be senescent cells. Senescent cells are characterized by cell cycle arrest and thus their G2M scores and S scores are expected to be low. We found that some cells in low G2M score and low S score have relatively low CFS percentiles (Methods, Figure 3c). However, after we updated the CFS scores using the significant new senescence markers (Figure 3d), or the combined markers (Figure 3e), most cells in this region have elevated CFS percentiles. This suggests that the new senescence markers and the combined markers have increased our ability to identify more senescent cells compared with only using the input markers.

### B-Lightning Identifies new T effector and memory markers in COVID-19 patients and revealed synchronized dynamics between effector and memory T cells

The pandemic of Coronavirus disease 2019 (COVID-19) has posed devastating effects to human health and society in the world^[18]^. Studying the immune system response to the SARS-COVID-2 viruses is critical to understanding the pathogenesis of COVID-19^[19,20]^ and gaining insights towards preventing the next pandemic. T cells are a major component of the adaptive immune system and play a critical role in the immune response against SARS-COVID-2 viruses. In this study, we sought to study the dynamics of T effector and memory cells in COVID-19 patients, and how the dynamics relate to the COVID-19 patient outcome.

We started from a dataset of a 74-year-old female patient of severe COVID-19^[15]^ with lung tissue CD4 and CD8 T cells extracted, and gene expression measured along the days during hospitalization (1, 3, 4, 5, 6, 7, 8 days post intubation) (Supplementary Figure 1a, 2a). We applied B-Lightning to enrich T cell effector and memory markers (Methods). After enrichment, for CD4 T cells with the combined markers (input markers and the newly identified markers), the median of memory CFS exhibits large increase in Day 4 and decrease in Day 5, and remains relatively low after Day 5. The median of effector CFS exhibits the opposite trend, showing large decrease in Day 4 and increase in Day 5, and remains relatively high after Day 5 (Figure 4a, b). After Day 5, the effector CD4 T cells expand again, possibly due to the uncontrolled SARS-COVID-2 virus infection for this severe symptomatic patient. The memory CD4 T cells keep at a high abundance. The result aligns with the previous study where researchers found that convalescent individuals following COVID-19 exhibited broad and strong memory T cell responses, especially in severe cases^[21]^. We checked the expression level of exhausted CD4 T cell markers: PDCD1, TIM3, LAG3, CTLA4 and ENTPD1 (CD39)^[22,23]^ among cells with high effector CFS on each day (effector CFS > 0.8), and found that the expression level of these markers increased on day 5 and afterwards (Figure 4c). This suggests that although the effector CD4 T cells expanded again, they were functionally exhausted. This is consistent with the previous COVID-19 study that shows the surviving T cells appeared functionally exhausted, with elevated levels of exhaustion markers like PD-1^[24]^.

After marker enrichment by B-Lightning, we observed a synchronized dynamic pattern between effector and memory CD4 T cells: as memory T cells expand, effector T cells decrease, and vice versa (Figure 4a, b). This pattern cannot be revealed only based on the input markers (Supplementary Figure 5a). We further checked the Pearson correlation between the effector and memory CFS. After B-lightning enrichment, the memory and effector CFS show significant negative correlations, even during day 5-7 where few cells were collected (Figure 4d). However, before the enrichment, we cannot observe negative correlations on any of the days (Figure 4e). To verify this observation, we applied the same combined marker set to the 66-year-old and 49-year-old COVID-19 patient datasets to calculate the memory and effector CFS. The synchronized dynamics and negative correlations across days are consistent with the 74-year-old patient dataset (Figure 5). Furthermore, B-lightning enrichment also reveals the synchronized patterns and significant negative correlations between memory and effect CFS for CD8 T cells on the most of the days (except for Day 5-7 of the 74-year-old CD8 dataset where too few cells were collected). Similarly, without B-lightning enrichment, the significant negative correlations are observed on very few days (Supplementary Figures 3, 4, 5, and 6). This suggests that B-Lightning helps to boost the signals of effector and memory functions and reveals the dynamic transition between effector and memory T cells.

We also annotated the cells with their memory and effect CFS. With input markers only, cells with high memory CFS and effector CFS are mixed together, introducing difficulties in interpretation (Supplementary Figure 1b,d Supplementary Figure 2b,d). With combined markers, cells with high memory CFS and effector CFS are well separated (Supplementary Figure 1c, e Supplementary Figure 2c, e). Thus, B-Lightning helps to boost the effector and memory signals to better differentiate the T cells.

## Discussion

B-Lightning uses two checking steps, DGE checking step and gene co-expression graph edge connectivity checking step, to identify marker genes. In the current algorithm, the first step is to screen out differential expressed genes, and the second step is to further filter with edge connectivity. Technically speaking, these two steps can be reversed. In practice, we found that the current order is more computationally efficient because estimating gene coexpression graph based on quantile association is usually more time-consuming than DGE analysis. However, in some cases, the gene co-expression graph might be more informative than DGE analysis, and we can consider reversing the order of these two steps.

Another main step in the B-Lightning is to compute gene enrichment score based on the current set of markers. We proposed CFS based on the over- and under-expressions of the marker genes. Alternative gene enrichment methods are available, such as GSVA^[9]^, scGSEA^[25]^, or AUCELL^[26]^. In practice, we found that CFS is robust and computationally efficient, and perform well as a step in the B-Lightning algorithm. However, if future alternative methods show better potential in accurately calling cells with marker genes, this step can be switched to other methods.

We suggest using verified or consensus markers as input markers for B-Lightning. These markers usually have large log fold change to differentiate relevant cells from other heterogeneous cohorts so that they can be easily verified in multiple studies. Thus, it is not surprising if B-Lightning identified markers have a smaller log fold change than the input markers. Thus, further verification of the genes can be based on comprehensive literature review and additional experimental validation.

## Conclusion

In this study, we developed a new method, B-Lightning, that “fishes out” new markers based on tan input set of existing known markers. B-Lightning starts with a few validated and highly trustworthy bait genes and combines DGE analysis check and gene co-expression edge connectivity check to identify new context-specific markers. Due to this double-checking process, B-lightning is more robust and accurate than competing methods, which usually use one checking criterion. Multiple dataset analysis has demonstrated that B-lightning performs well in identifying new markers to boost the cell differentiation signals and reveal complex cell heterogeneity in single-cell data. Especially, B-Lightning can robustly identify markers with high precision, and thus low false discovery rate. Identifying additional markers increase the SOH signal-to-noise ratio and helps to differentiate the corresponding cell subpopulations of interest from other heterogeneous cohorts. This helps our downstream analysis to better understand the biological processes and states of interest.

## Key Points

- B-Lightning is a highly accurate and robust method for identifying marker genes and corresponding cell subpopulations linked to a specific source of heterogeneity (SOH) in single-cell data.
- By setting two thresholds based on gene differential expression patterns and gene coexpression network connectivity, B-Lightning effectively enriches SOH-specific marker genes while significantly reducing false discoveries.
- B-Lightning successfully identified novel potential senescence markers in lung ciliated cells and pinpointed a senescence-enriched subpopulation within these cells.
- B-Lightning also enriched T-cell effector and memory markers, uncovering a previously unknown synchronized dynamic pattern between these two T-cell subpopulations—a pattern that was not evident before B-Lightning’s marker enrichment.

## Data availability

The IPF datasets are collected from study GSE135893^[27]^. The 74-year-old female, 66-year-old female, and 49-year-old male airway longitudinal CD4/CD8 dataset are available at https://cellxgene.cziscience.com/collections/29f92179-ca10-4309-a32b-d383d80347c1. COVID-19 airway CD4/CD8 dataset is available at https://figshare.com/articles/dataset/COVID-19_severity_correlates_with_airway_epithelium-immune_cell_interactions_identified_by_single-cell_analysis/12436517.

## Code availability

The R package B-Lightning v0.1 is available at https://github.com/yirenshao/B-Lightning. The codes for generating the results shown in this paper are available at https://github.com/jichunxie/Blightning_manu/.

## Acknowledgement

The authors thank the Duke Center for Human Systems Immunology (CHSI) for providing the computation resources.

## Funding

YS, QG, AN, and CC’s research was supported by NIH Common Fund, through the Office of Strategic Coordination/Office of the NIH Director under awards, 1U54AG075936 from the National Institute on Aging. LW’s research was supported by Duke University. DL’s research was supported by National Institute of Environmental Health Sciences with grant number NIH 1R21ES032159 and National Institute on Aging with grant number NIH 1U54AG075931. Q-JL’s research was supported by core research grants provided to the IMCB and SIgN by the BMRC, A*STAR and National Research Foundation (NRF) Singapore under the NRF Investigatorship NRFI09-0016. And JX’s research was supported by National Human Genome Research Institute with the grant number NIH 1R01HG012555 and National Instite on Aging with the grant number 1U54AG075936. The content is solely the responsibility of the authors and does not necessarily represent the official views of the National Institutes of Health.

## Conflict of Interest Statement

The authors declare no competing interests.

## Notes

### Competing Interest Statement

The authors have declared no competing interest.

