## supplementary note for "Marker gene fishing for single-cell data with complex heterogeneity"

### Supplementary Note 1: Quantile association statistics $S_{ij}$ and the connectivity testing

We followed the same procedure as SifiNet[1] to use quantile associations to measure gene co-expression. The details are described below.

Let $Y_{ic}$ be the feature values of gene $i$ in cell $c$. For scRNA-seq data, the gene feature value could be the gene UMI count. Based on the gene feature values, we construct a gene co-expression network among all the genes. Each node represents a gene, and each edge represents the co-expression between two genes.

Let $s_c$ be cell $c$'s library size (such as total read counts). We use quantile associations to define gene co-expressions.

$$D_{ij} = \frac{1}{N}\sum_{c} P(Y_{ic} \leq q_{ic}, Y_{jc} \leq q_{jc} \mid s_c) - \tau_i\tau_j,$$

where $q_{ic}$ and $q_{jc}$ are the $\tau_i$-th and $\tau_j$-th conditional quantiles of gene $i$ and gene $j$'s expressions. $s_c$ can be further extended to include other cell-specific covariates. In the gene-co-expression network, if $D_{ij} \neq 0$, there is an edge between nodes $i$ and $j$.

Here, we establish a standardized estimator $S_{ij}$ for the co-expression $D_{ij}$:

$$S_{ij} = \frac{\sum_{c=1}^{N}[I(Y_{ic} \leq \hat{q}_{ic}, Y_{jc} \leq \hat{q}_{jc}) - \hat{\tau}_i\hat{\tau}_j]}{\sqrt{N\hat{\tau}_i(1-\hat{\tau}_i)\hat{\tau}_j(1-\hat{\tau}_j)}},$$

where $\hat{q}_{ic}$ is the estimated cell specific $\tau_i$-th conditional quantile of gene $i$, and $\hat{\tau}_i = \sum_{c=1}^{N} I(Y_{ic} \leq \hat{q}_{ic})/N$. By default, we use the quantile regression model $q_{ic} = \alpha_{i0} + \alpha_{i1}s_c$ to estimate $\hat{q}_{ic}$, where $\tau_i = (1+\hat{p}_{i0})/2$ and $\hat{p}_{i0} = \sum_{c=1}^{N} I(Y_{ic} = 0)/N$ is the proportion of zero observations of gene $i$. In this default setting, the estimated $\hat{q}_{ic}$ corresponds to the 50%-quantile among the non-zero observations of gene $i$ given cell library size. However, when gene $i$ has very low expression (when the median of $Y_{ic}$ is equal to 0 or $Y_{ic}$ are either 0 or 1), we set $\hat{q}_{ic} = 0$ for stability and robustness consideration.

Assume $Y_{ic}$ and $Y_{jc}$ are independent given a specific cell $c$. Our previous work[1] shows that if gene $i$ or gene $j$ is not a feature gene, then $D_{ij} = 0$, *i.e.*, these two genes are independent. Thus, its standardized measure $S_{ij}$ is used as a surrogate to measure if gene $j$ is also a feature gene, given gene $i$ is a feature gene.

To formalize the test, we extract highly variable genes from the scRNA-seq dataset using the recommended procedure by the function `FindVariableFeatures` in Seurate V3[2]. Denote the highly variable gene set by $\mathcal{H}$. For any $k \in \mathcal{H}$, we calculate $S_{ik}$ and use all $S_{ik}$ to form a benchmark null distribution to test the significance of $S_{ij}$: We claim gene $i$ and gene $j$ are connected only if $S_{ij}$ exceeds the 95% quantile of $S_{ik}$ for all $k \in \mathcal{H}$.

#### Supplementary Figures

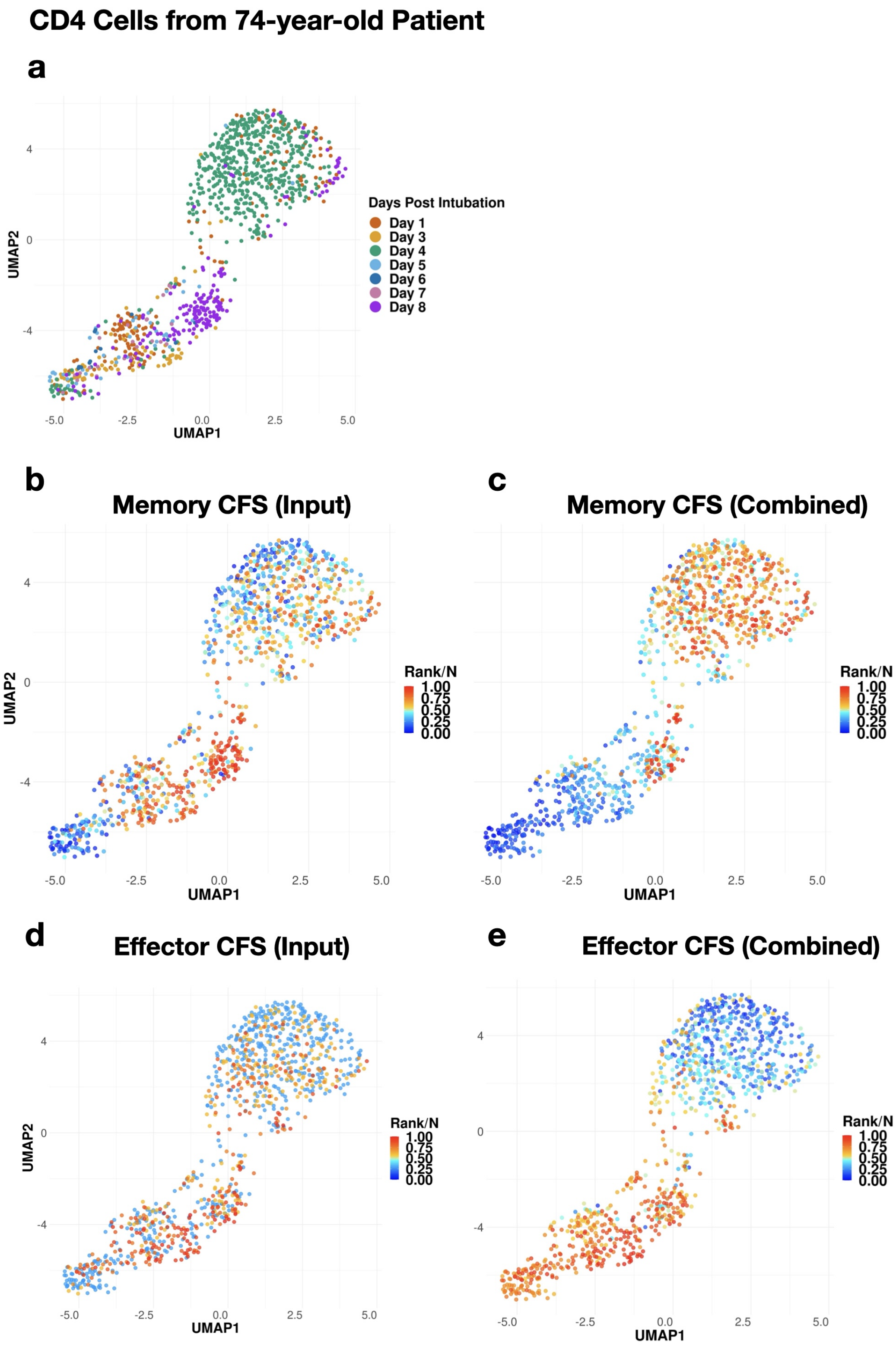


Supplementary Figure 1: **UMAP of airway CD4 cells from a 74-year-old female.** (a) Cells colored by days post intubation. (b) Cells colored by the ranking percentile of the memory CFS calculated by input markers. Cells with larger CFS are redder. (c) Cells colored by the ranking percentile of the memory CFS calculated by the combined markers. (d) Cells colored by the ranking percentile of the effector CFS calculated by input markers. (e) Cells colored by the ranking percentile of the effector CFS calculated by the combined markers.

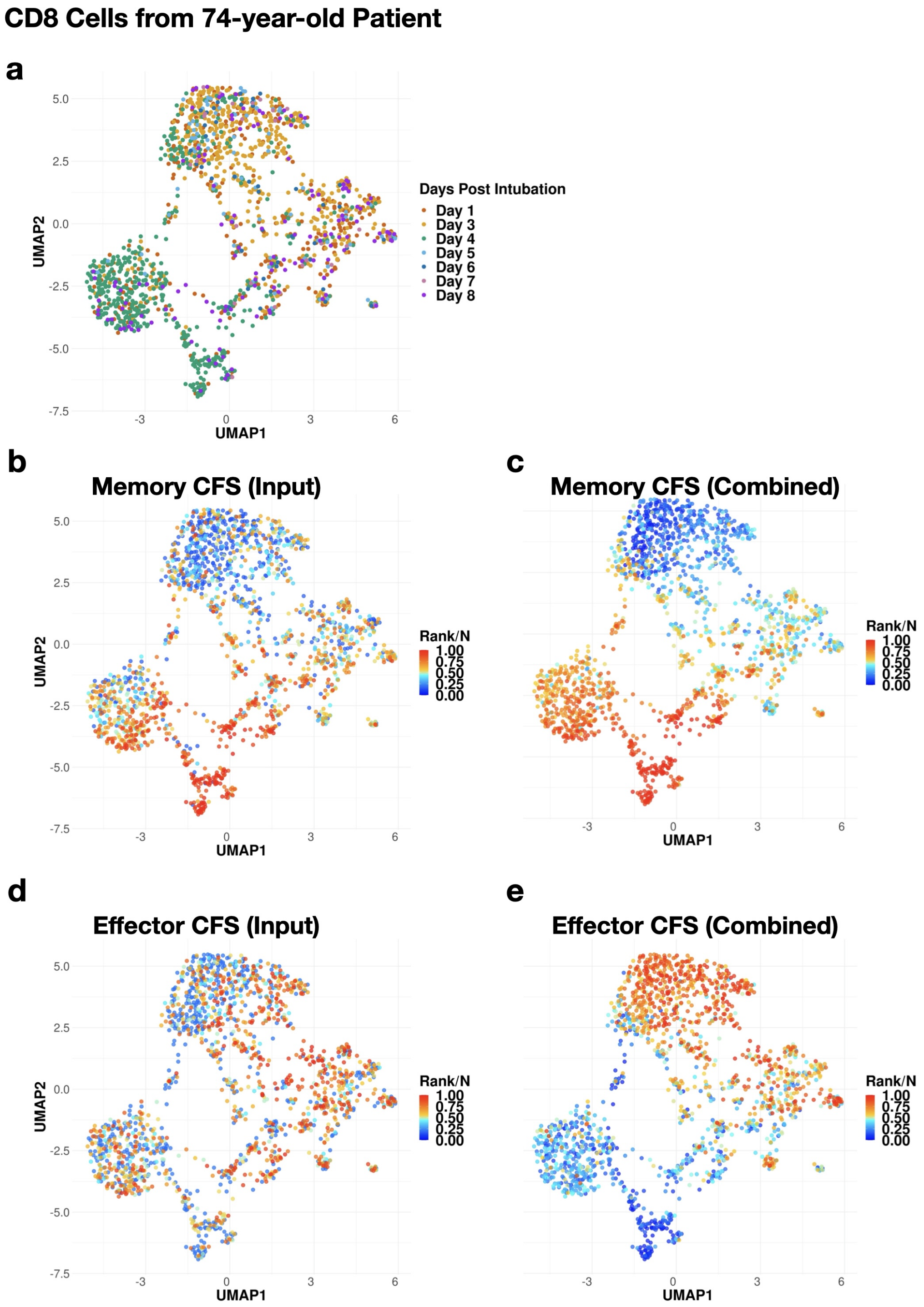


Supplementary Figure 2: **UMAP of airway CD8 cells from a 74-year-old female.** (a) Cells colored by days post intubation. (b) Cells colored by the ranking percentile of the memory CFS calculated by input markers. Cells with larger CFS are redder. (c) Cells colored by the ranking percentile of the memory CFS calculated by the combined markers. (d) Cells colored by the ranking percentileof the effector CFS calculated by input markers. (e) Cells colored by the ranking percentile of the effector CFS calculated by the combined markers.

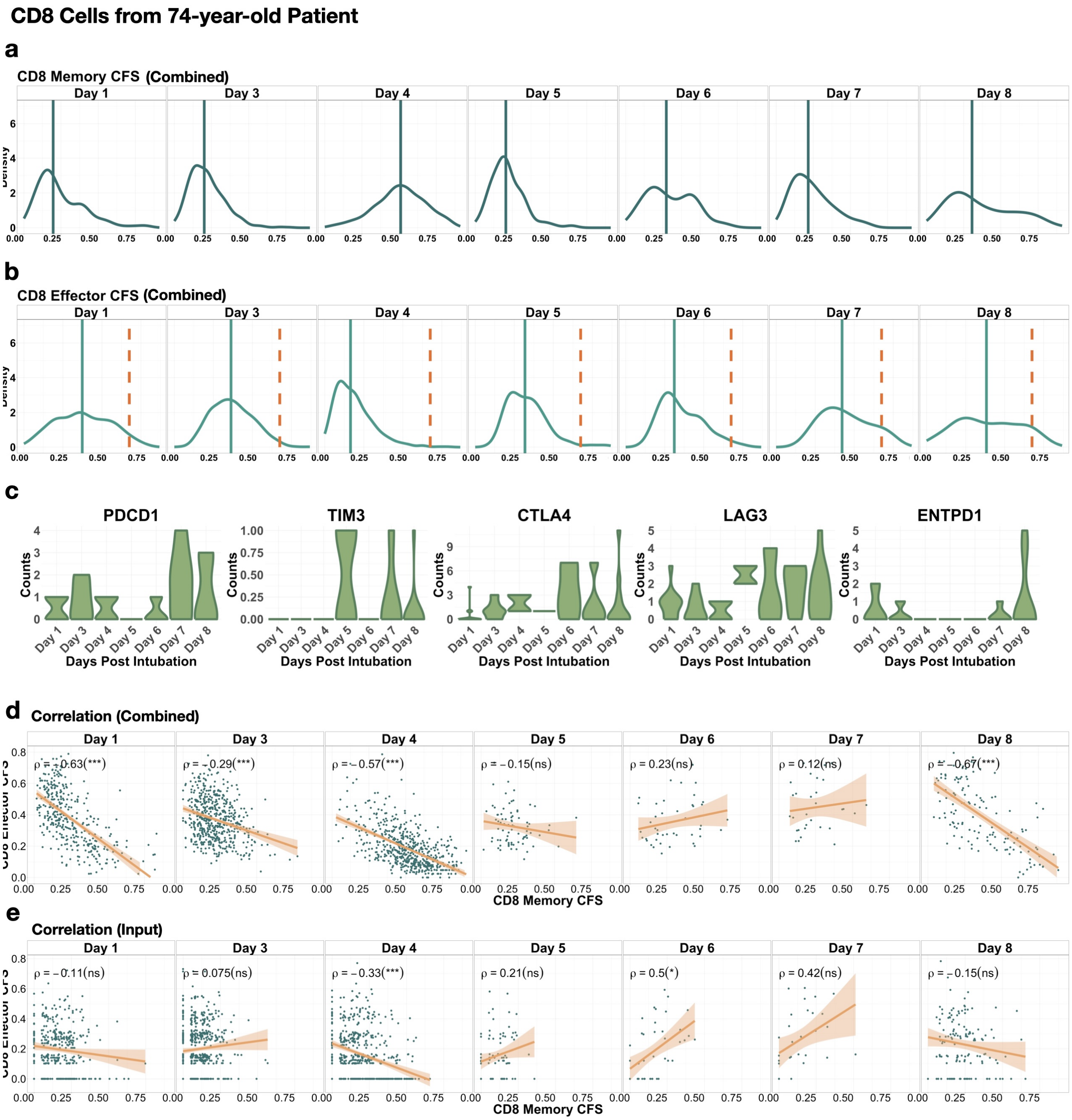


Supplementary Figure 3: **Density plots of CD8 CFS.** (a) The CD8 memory CFS. (b) The CD8 effector CFS. CFS was calculated by the combined markers. The vertical solid line refers to the median of the combined marker CFS. The orange dashed line refers to the combined marker effector CFS threshold (0.675) used to select cells with high effector CFS. (c) **Expression counts of CD8 cell exhaustion markers.** Vioin plots of T cell exhaustion markers PDCD1, TIM3, CTLA4, LAG3 and ENTP1 (CD39) of effector CD8 T cells (with combined marker effector CFS $> 0.675$) on days post intubation. **Scatter Plots of CD8 memory CFS vs. effector CFS.** (d) The combined marker memory CFS is the x-axis, and the combined marker effector CFS is the y-axis. (e) The input marker memory CFS is the x-axis, and the input marker effector CFS is the y-axis. $\rho$ stands for the Pearson correlation between the memory and effector markers. Asterisks/ns in the parentheses represent the significance level of the correlation between senescent and non-senescent cells, $*$: p-value $\leq 10^{-2}$, $**$: p-value $\leq 10^{-4}$, $***$: p-value $\leq 10^{-6}$, ns: non-significant. Each subplot extracted the cells collected on a specific day after intubation.

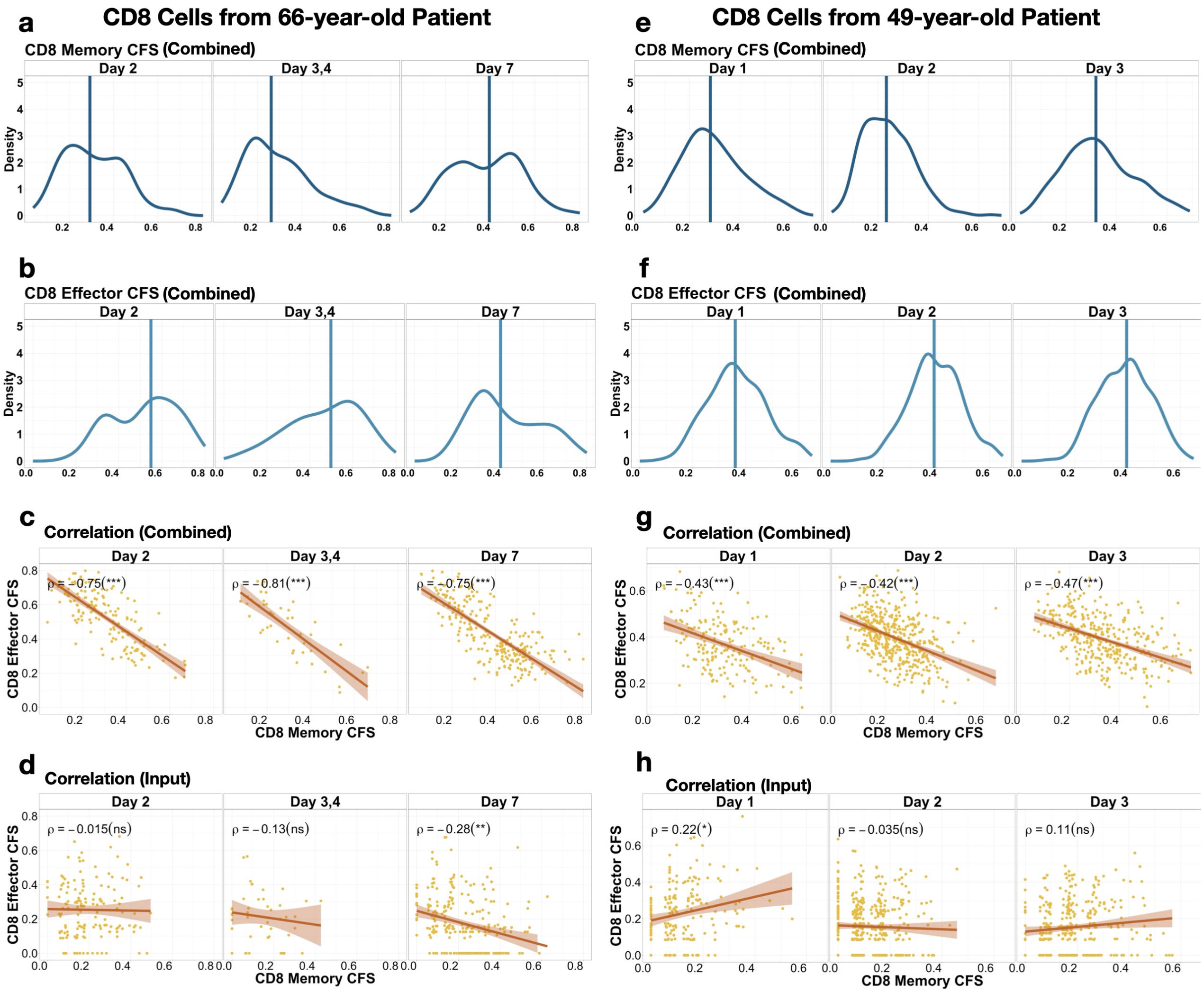


Supplementary Figure 4: **Density plots of combined marker CFS for CD8 cells from the 66-year-old and 49-year-old Patients.** (a,e) The combined marker CD8 memory CFS. (b,f) The combined marker CD4 effector CFS. The vertical solid line refers to the median of the combined marker CFS. **Scatter Plots of CD8 memory CFS vs. effector CFS.** (c,g) The combined marker memory CFS is the x-axis, and the combined marker effector CFS is the y-axis. (d,h) The input marker memory CFS is the x-axis, and the input marker effector CFS is the y-axis. $\rho$ stands for the Pearson correlation between the memory and effector markers. Asterisks/ns in the parentheses represent the significance level of the correlation between senescent and non-senescent cells, $*$: p-value $\leq 10^{-2}$, $**$: p-value $\leq 10^{-4}$, $***$: p-value $\leq 10^{-6}$, ns: non-significant. Each subplot extracted the cells collected on a specific day after intubation.

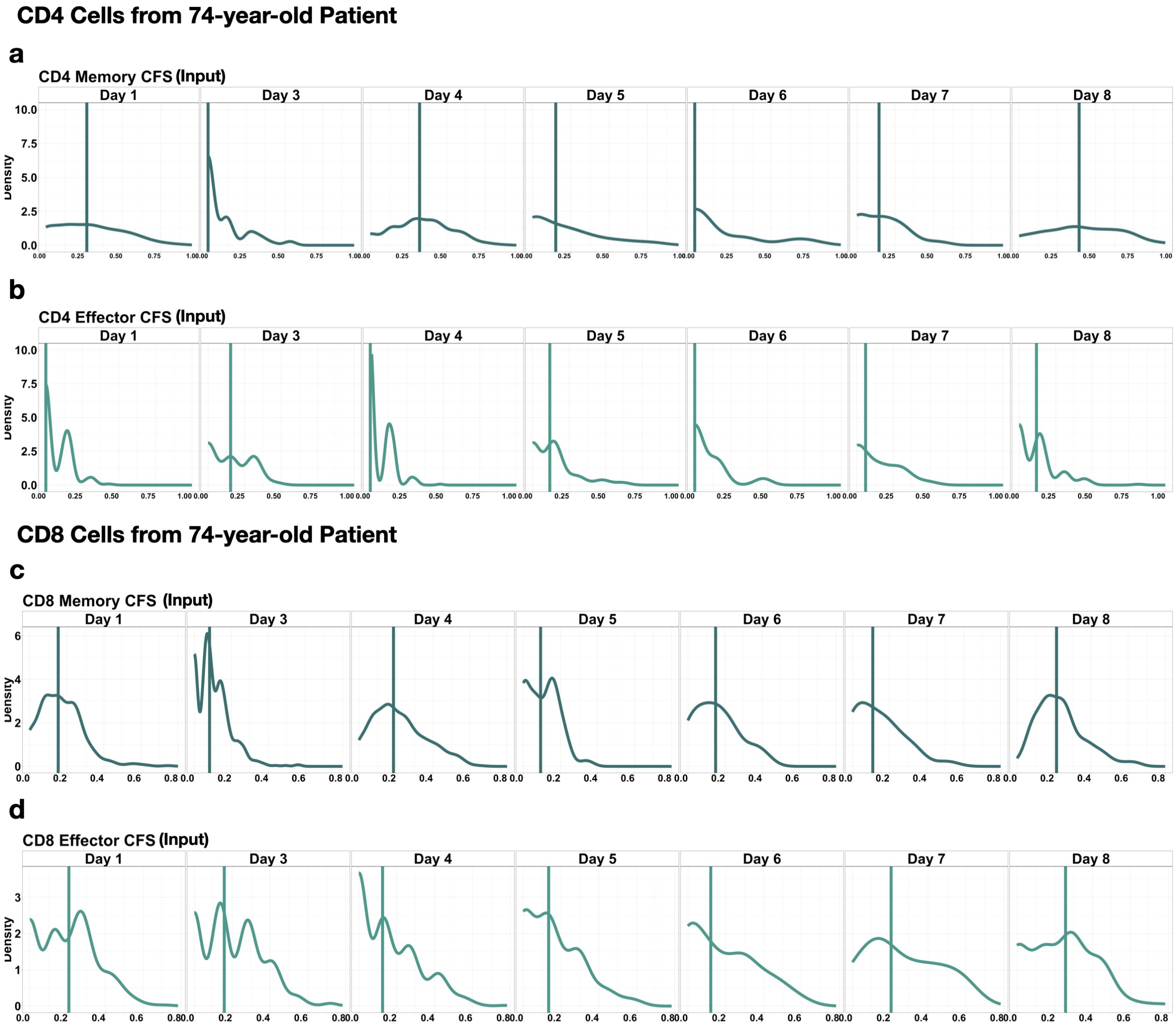


Supplementary Figure 5: **Density plots of input marker CFS of the 74-year-old Patient.** (a) The input marker CD4 memory CFS; (b) the input marker CD4 effector CFS; (c) the input marker CD8 memory CFS; (d) the input marker CD8 effector CFS.

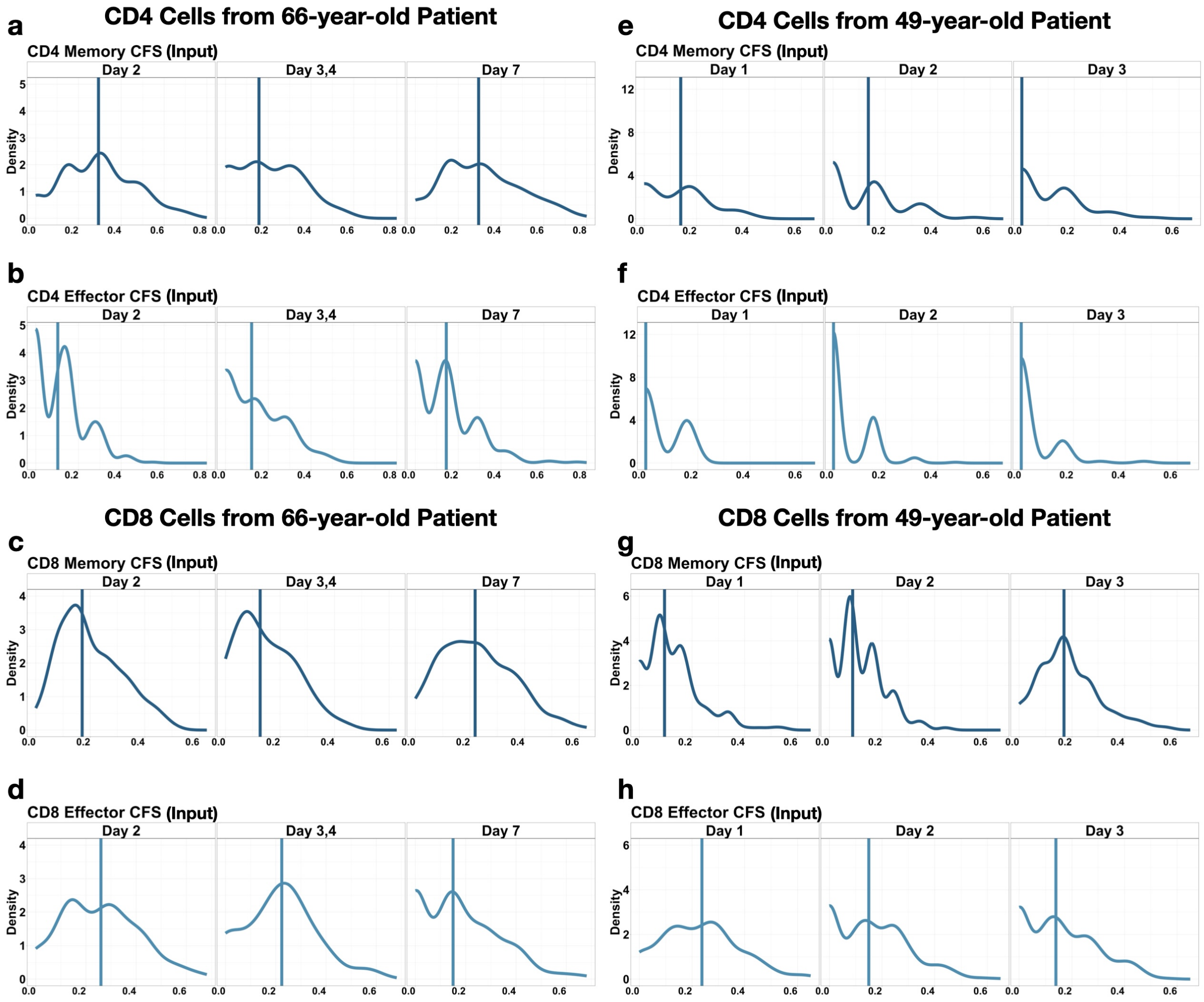


Supplementary Figure 6: **Density plots of input marker CFS of the 66-year-old and 49-year-old Patients** (a,e) The input marker CD4 memory CFS; (b,f) the input marker CD4 effector CFS; (c,g) the input marker CD8 memory CFS; (d,h) The input marker CD8 effector CFS.
